# Multiple lineages of *Monkeypox virus* detected in the United States, 2021- 2022

**DOI:** 10.1101/2022.06.10.495526

**Authors:** Crystal M. Gigante, Bette Korber, Matthew H. Seabolt, Kimberly Wilkins, Whitni Davidson, Agam K. Rao, Hui Zhao, Christine M. Hughes, Faisal Minhaj, Michelle A. Waltenburg, James Theiler, Sandra Smole, Glen R. Gallagher, David Blythe, Robert Myers, Joann Schulte, Joey Stringer, Philip Lee, Rafael M. Mendoza, LaToya A. Griffin-Thomas, Jenny Crain, Jade Murray, Annette Atkinson, Anthony H. Gonzalez, June Nash, Dhwani Batra, Inger Damon, Jennifer McQuiston, Christina L. Hutson, Andrea M. McCollum, Yu Li

**Author notes:** The findings and conclusions in this report are those of the author(s) and do not necessarily represent the views of the Centers for Disease Control and Prevention or Los Alamos National Laboratory.

## Abstract

Monkeypox is a viral zoonotic disease endemic in Central and West Africa. In May 2022, dozens of non-endemic countries reported hundreds of monkeypox cases, most with no epidemiological link to Africa. We identified two lineages of *Monkeypox virus* (MPXV) among nine 2021 and 2022 U.S. monkeypox cases. A 2021 case was highly similar to the 2022 MPXV outbreak variant, suggesting a common ancestor. Analysis of mutations among these two lineages revealed an extreme preference for GA-to-AA mutations indicative of APOBEC3 cytosine deaminase activity that was shared among West African MPXV since 2017 but absent from Congo Basin lineages. Poxviruses are not thought to be subject to APOBEC3 editing; however, these findings suggest APOBEC3 activity has been recurrent and dominant in recent West African MPXV evolution.

---

Monkeypox is a viral zoonotic disease caused by *Monkeypox virus* (MPXV) endemic in West and Central Africa. There have been several reported cases of travel-associated monkeypox in non-African countries in recent years. In 2003, an outbreak of monkeypox in the United States (U.S.) was linked to imported African small mammals (*1*). In 2017, the largest monkeypox outbreak in western Africa occurred in Nigeria after decades of no identified cases (*2*), and during 2018 to 2021, eight cases were imported from Nigeria to non-endemic countries(*2–8*). Of these, four occurred in the United Kingdom, and one each occurred in Israel and Singapore (*4–6, 8, 9*). In 2021, there were two U.S. monkeypox cases in travelers from Nigeria, one in Maryland and the other in Texas (*3, 7, 10*). In May of 2022, this pattern of monkeypox cases being reported in persons who had travelled in Nigeria shifted, and multiple countries reported monkeypox among persons who had not travelled to Nigeria or other monkeypox-endemic countries(*11*). As of June 8, 2022, 1200 cases of monkeypox cases were reported in 29 non-endemic countries, most with no epidemiolocal link to Africa. As of June 8, forty cases have been confirmed in the U.S.(*12*). The origin of this outbreak and the seemingly rapid transmission is currently under investigation. Viral genomic sequencing can be used to determine the similarity between viruses and suggest possible links between cases, origins of infection, and transmission dynamics when combined with epidemiological information.

Comparison of the U.S. MPXV sequences from May 2022 (ON563414.3, ON674051, ON675438, ON676703 – ON676708) revealed two distinct lineages (Figure 1, red and blue). The majority (5/7) of 2022 U.S. sequences formed a monophyletic clade with 2022 MPXV sequences from Europe (Figure 1, red), with most genomes containing 0 – 1 nucleotide changes in non-repeat regions (Figure 2, see methods for genome comparison). This clade will be referred to as the current predominant 2022 MPXV outbreak clade, although increased surveillance and sequencing may reveal a different predominant outbreak strain in the future. MPXV from a 2021 travel-associated case from Nigeria to Maryland (USA_2021_MD) displayed high similarity to the predominant 2022 MPXV outbreak sequences, with approximately 13 nucleotide differences (Figures 1, pink, and 2, Table 1). The USA_2021_MD and 2022 outbreak sequences contained many shared mutations that separated them from MPXV sequences from Nigeria and other travel-associated cases from 2017 – 2019 (Supplemental Table 1).

**Figure 1.**
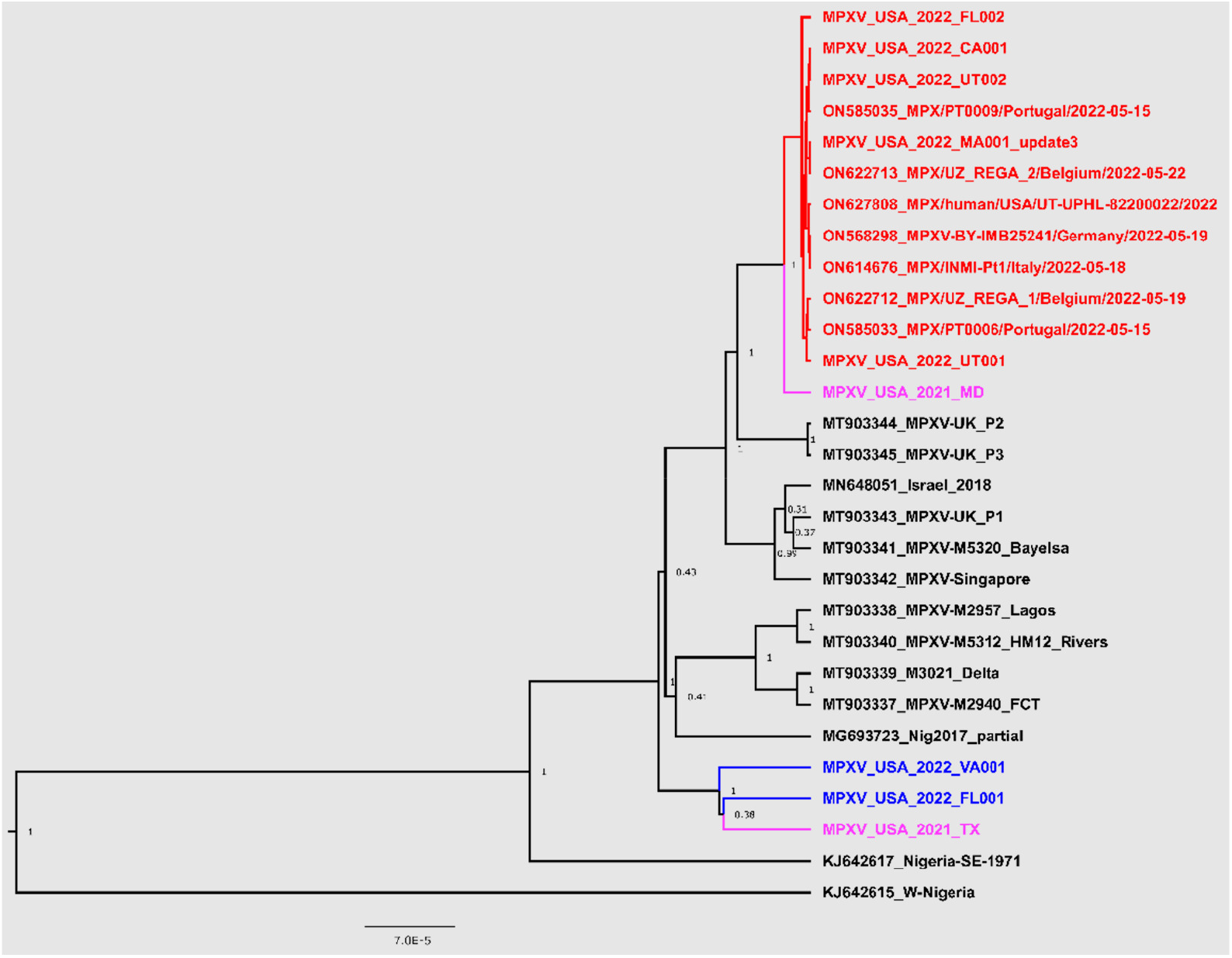
Phylogenetic analysis of West African MPXV genome sequences from the 2022 outbreak, recent travel-associated cases (2018 – 2021), and MPXV from Nigeria. Phylogenetic analysis was performed in BEAST v1.8.3 using HKY+G model and constant coalescent prior. Whole genome sequence alignment was generated using MAFFT v.7.450 (*28*) followed by removal of alignment columns containing gaps. Scale bar is in substitutions per site; posterior support values are shown at branch points.

**Figure 2.**
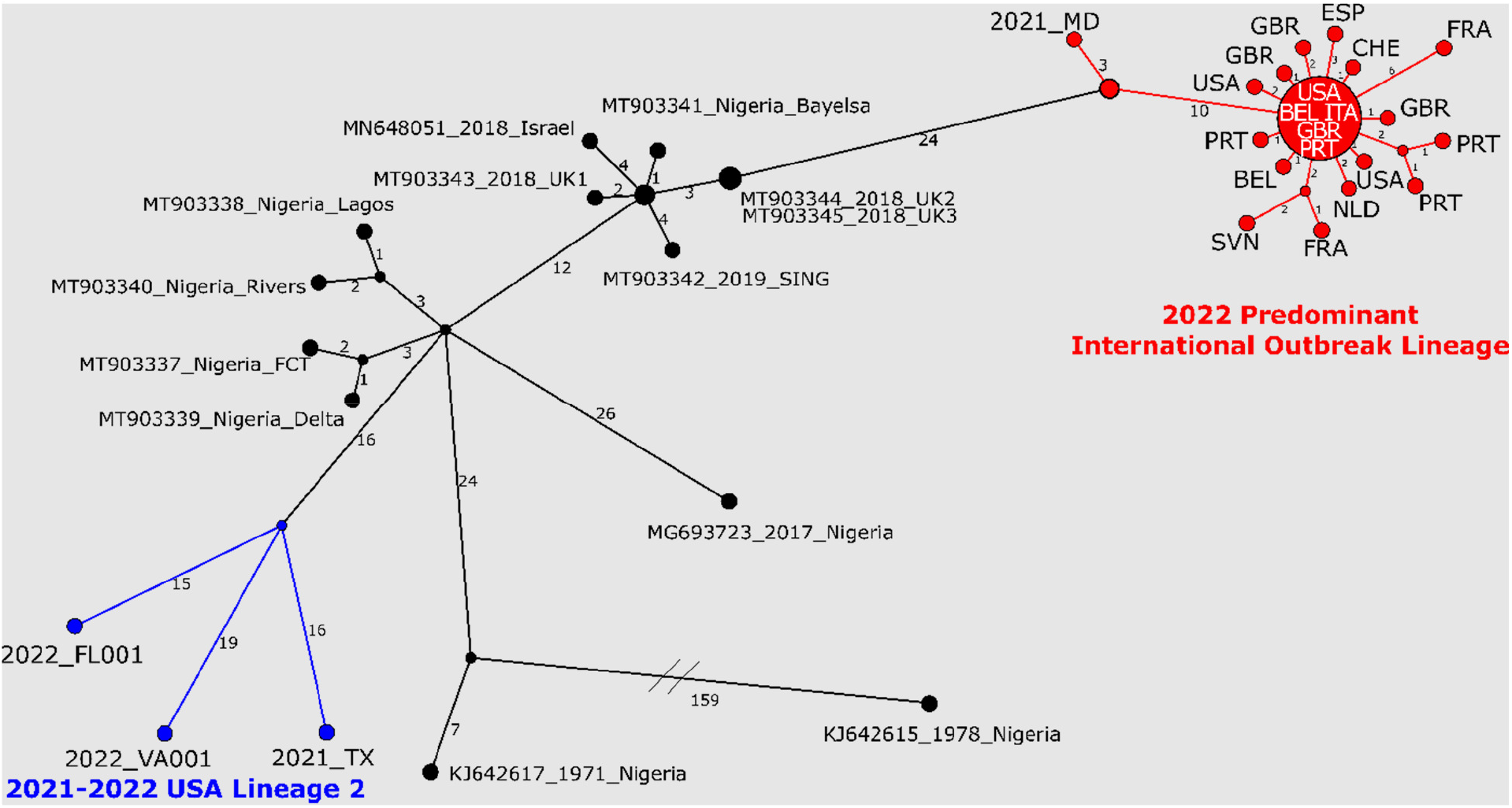
Nucleotide changes among West African MPXV genome sequences from 2022 outbreak, recent travel-associated cases (2018 – 2021), and cases from Nigeria. The predominant 2022 MPXV outbreak lineage is highlighted in red; 2021 TX, 2022 FL002, and 2022 VA001 are shown in blue. 2022 predominant MPXV outbreak cluster sequences are represented using the three letter code for the country; the large node at the center of the predominant 2022 MPXV outbreak cluster represents 13 identical sequences (sequences used can be found in Supplemental Table 4). Haplotype network analysis was generated with PopArt using the Median Joining method. Sequence differences between nodes are indicated by the numbers on the branches. Unlabeled nodes represent hypothetical common ancestors, lines connecting nodes show the number of sequence differences and do not represent direct links between cases. Sequence alignment was generated by whole genome alignment using MAFFT v.7.450 (*28*) followed by removal of alignment columns containing gaps or ambiguities.

**Table 1.** Nucleotide changes of between USA_2021_MD and USA_2022_MA001. MA001 (ON563414.3) was used as a representative genome for the predominant 2022 outbreak cluster. Gene homologs in *Vaccinia virus* strain Copenhagen are given for each gene as well as notes on proposed functions taken from annotation of MT903345. Additional information including mutations in non-coding regions can be found in Supplemental Table 1.

| Position | 2021_MD | 2022_MA001 | TYPE | EFFECT | NT CHANGE | AA CHANGE | VACV-Cop | Note |
| --- | --- | --- | --- | --- | --- | --- | --- | --- |
| 3120 | G | A | snp | synonymous | 1497 C > T | - | C19L | Ankyrin |
| 39148 | C | T | snp | missense | 1057 G > A | Glu353Lys | F13L | EEV membrane glycoprotein |
| 73248 | G | A | snp | missense | 262 G > A | Asp88Asn | G8R | VLTF-1 (late transcription factor 1) |
| 74214 | G | A | snp | missense | 426 G > A | Met142Ile | G9R | Entry/fusion complex component |
| 77392 | G | A | snp | missense | 484 G > A | Glu162Lys | L4R | ss/dsDNA binding protein (VP8) |
| 79357 | T | C | snp | synonymous | 264 C > T | - | J3R | Cap-specific mRNA-methyltransferase |
| 84596 | C | T | snp | synonymous | 3462 C > T | - | J6R | RNA polymerase subunit (RPO147) |
| 86988 | T | C | snp | synonymous | 81 G > A | - | H3L | IMV surface membrane protein |
| 150480 | C | T | snp | missense | 661 C > T | His221Tyr | A46R | IL-1/TLR signaling inhibitor |
| 169732 | - | ACAGAT | del | deletion | 100 - 105 del | del:Thr34,Asp35 | B11R | Hypothetical protein |
| 170273 | G | A | snp | synonymous | 225 G > A | - | B12R | Ser/Thr Kinase |
| 183534 | C | T | snp | missense | 2164 C > T | Pro722Ser | B21R | Surface glycoprotein |
| 194114 | C | T | snp | synonymous | 1497 C > T | - | C19L | Ankyrin |

Two 2022 U.S. MPXV sequences, USA_2022_FL001 and USA_2022_VA001, were polyphyletic to other 2022 MPXV sequences from the U.S. and Europe (Figure 1, blue). Each genome contained approximately 80 nucleotide changes relative to the other 2022 MPXV sequences (Figure 2), indicating these were introductions unrelated to the other May 2022 U.S. cases. Additionally, these two sequences were most similar to MPXV from a 2021 traveler from Nigeria to Texas. The 2022 MPXV from FL001 and VA001, however, displayed approximately 30 unique nucleotide differences from each other and USA_2021_TX (Figure 2, Supplemental Table 2), suggesting the cases were not directly linked.

Real-time PCR testing of USA_2021_MD and the five 2022 MPXV outbreak samples revealed a lower sensitivity (higher cycle threshold (Ct) value) when compared to published *Orthopoxvirus* generic OPX3 real-time PCR assay (*13*) compared to Monkeypox West African-specific real-time PCR assay (*14*) (supplemental Table 3, when performed as described in Methods). An average Ct delay of 6.88 (ranging from 5.3 to 8.3) in the OPX3 assay compared to West African specific assay was observed in the USA_2021_MD and five 2022 MPXV outbreak cluster samples. By contrast, the OPX3 and West African-specific assay produced similar Ct values for USA_2022_FL001, USA_2022_VA001, or USA_2021_TX samples (average difference of −0.78 Ct, ranging from −1.4 to 0.5). Sequence examination revealed a SNP in the reverse primer binding site for the *Orthopoxvirus* OPX3 real-time PCR assay (DNA polymerase gene, VACV-Cop E9L, 322 C to T) in USA_2021_MD and the five 2022 outbreak cluster sequences that is absent from other West African MXPV sequences. This SNP is conserved in 2022 MPXV sequences from Europe.

When comparing the 2022 outbreak MPXV sequences to other MPVX sequences, it was evident that many of the observed mutations in the 2022 sequences were 5’ GA-to-AA, which is indicative of Apolipoprotein B mRNA Editing Catalytic Polypeptide-like3 (APOBEC3) activity (*15*). We performed a comparison of G-to-A changes within the APOBEC3 motif (5’ GA-to-AA) to other G-to-A changes across West African and Congo Basin MPXV lineages. We found no evidence for an enrichment of G-to-A mutations in West African MPXV clades prior to 2017 or Congo Basin MPXV clades (Fig. 3A,B, Supplemental Figure S1). In contrast, we found a profound enrichment of G-to-A mutations in the more recently sampled West African MPXV sequences (2017 to 2022); we found 160 G-to-A mutations in an APOBEC3 motif, 9 G-to-A mutations not in an APOBEC3 motif, and only 7 mutations that were not G-to-A. The vast majority of the APOBEC3 context G-to-A mutations were specifically GA-to-AA (156 out of 160, 96%), indicating that these mutations were not generated by APOBEC3G (produces GG-to-AG changes), but rather by other APOBEC3 subfamily members (*16, 17*).

**Figure 3.**
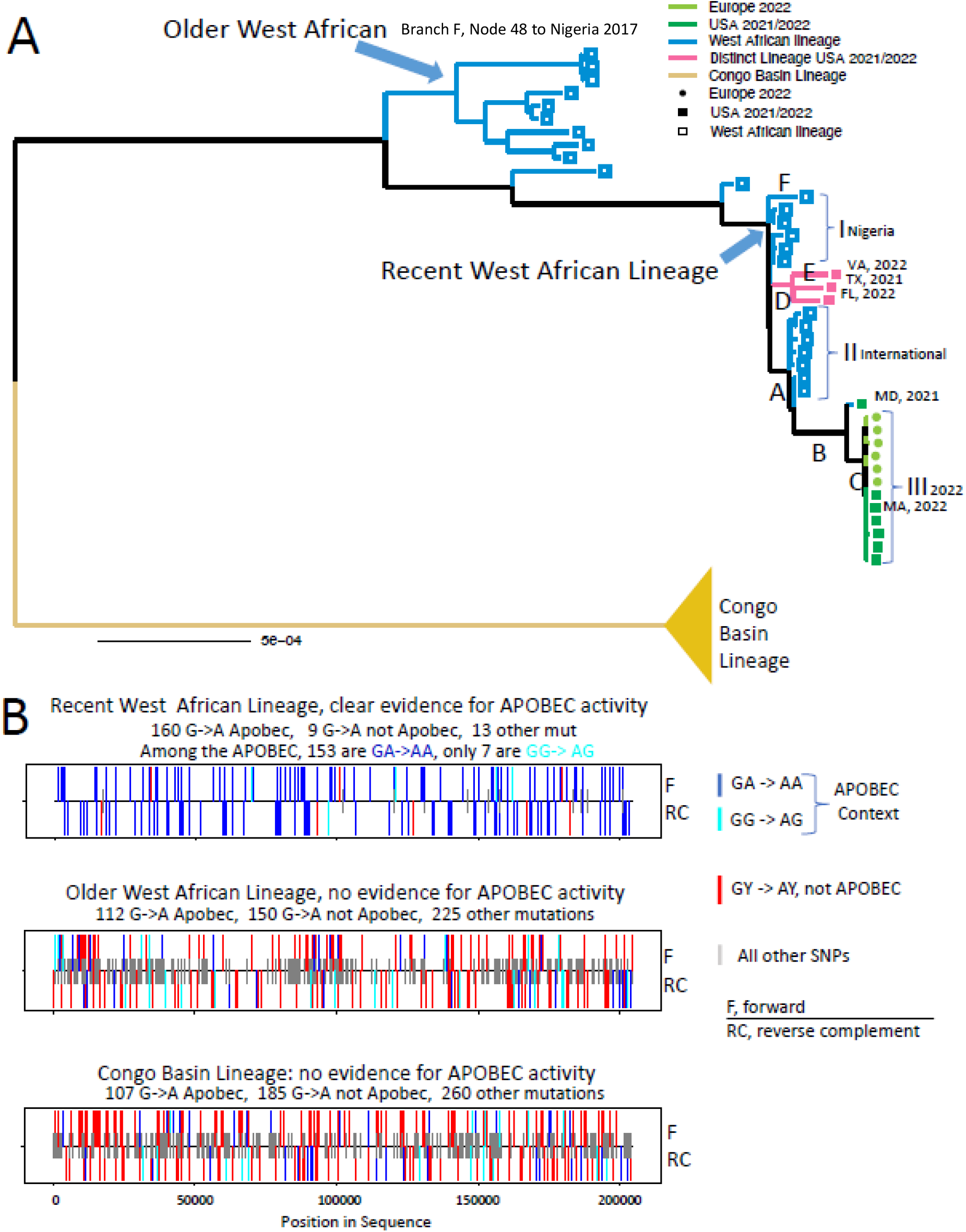
Analysis of APOBEC3 motif mutations in the West African MPXV lineage. A. Maximum Likelihood phylogenetic tree using IQ-TREE (*29*). A detailed tree, with taxa names is included in Fig. S1. The scale bar indicates the number of substitutions per sequence site. Letters A-E indicate branches that were each independently significantly enriched for G-A mutations embedded in an APOBEC3 motif. Clusters I, II, and III represent clusters of highly related sequences that have almost exclusively G-A mutations embedded in an APOBEC3 motif relative their most recent common ancestor. B. Mutational patterns found among the three major lineages in the phylogeny above. All unique mutations in a clade relative to the most recent common ancestor of that clade are compressed onto a single sequence, and the class of each mutation is show on either the forward or reverse complement strand. Blue bars indicate GA-to-AA (an APOBEC3 motif used by many APOBEC3 subfamily members, but not used by APOBEC3G), cyan indicates GG-to-AG (APOBEC3G motif), and red indicates GY to AY (non-APOBEC3 motif); shorter gray lines indicate all other single nucleotide polymorphism (SNP) mutations. Bars above the horizontal line indicates forward strand, below the line indicates reverse strand.

We then explored the relative abundance of G-to-A mutations in an APOBEC3 motif along the full evolutionary trajectory from the common ancestor of 2017 Nigeria MPXV to the two MPXV lineages identified in 2021 - 2022 U.S. cases (Figures 1 and 3, Supplemental Figure 1). The full path from the estimated ancestor of 2017 – 2022 West African MPXV sequences to the recent 2022 outbreak, represented by MPXV_USA_2022_MA001_V3 (ON563414.3) was significantly enriched for APOBEC3 vs non-APOBEC3 motif G-to-A mutations (p < 0.000001). Each branch along the way was also significantly enriched for APOBEC3 G-to-A mutations, suggesting sequential rounds of mutation in distinct individuals gave rise to the high proportions of GA-to-AA APOBEC3 motif mutations ultimately shared by the 2022 outbreak viruses (Figure 3, Supplemental Figures 2 and 3). The 2021 Maryland MPXV (MPXV_USA_2021_MD) captured an intermediate step along the way. Similarly, the path to the second MPXV lineage in the U.S., represented by the sequence MPXV_USA_2021_TX, also was highly enriched for APOBEC3 motif mutations (p = 0.000001), as were individual branches and each of the terminal branches (Figure 3, Supplemental Figures 2 and 3). Finally, significant enrichment of APOBEC3 motif G-to-A mutations were found on the branch labeled F in Figure 3, leading to a Nigerian sequenced sampled in 2017 (Supplemental Figures 2 and 3).

Finally, we summarized all unique mutations found in three clusters of highly related West African MPXV viruses, labeled shown in Figure 3, relative to their most recent common ancestor: Cluster I, recent samples from Nigeria (2017); Cluster II, recent non-Nigerian samples (2017-2019); and Cluster III, U.S. and European samples from May 2022. Though there are only a small number of mutations leading from the shared ancestor to any one sequence, the overwhelming preponderance of mutations across each cluster were GA-to-AA substitutions in an APOBEC3 context, with significant enrichment across each group (Supplemental Figures 4-6). This suggests that APOBEC3 is acting at low levels in many hosts, accumulating mutations over series of infections.

All publicly available MPXV genomes from the 2022 Monkeypox outbreak to date belong to West African MPXV lineage, which may cause less severe disease and have a lower case fatality than Congo Basin MPXV clade (*18–20*). Genomes published during the 2022 monkeypox outbreak share a common ancestor with MPXV from Nigeria; however, sequences from surrounding countries are limited and most of our understanding of these relationships comes from viruses linked to or identified in Nigeria. The high similarity among the current predominant European and U.S. 2022 MPXV strain is typical of what has been reported for samples from the same transmission chain (0.4 to 1.5 SNPs per genome) (*4*). Most of the unique mutations observed in the predominant 2022 MPXV lineage were shared with USA_2021_MD, further indicating that any evolutionary force leading to these changes most likely preceded the 2022 outbreak. However, there were sufficient differences (>30 SNPs) among the 2021_TX, 2022, 2022_VA001 and 2022_FL001 sequences to suggest they are likely not part of the same transmission chain. Furthermore, each of the three cases had traveled to different countries, so at this time it is unclear how widely dispersed this lineage of MPXV is until more sequences are available.

From 2017 to 2022, 588 cases of monkeypox were reported in Nigeria, with only 15 confirmed cases in 2022 as of April 30 (*21*). The current number of confirmed cases around the world in persons with no travel history to endemic countries and the seemingly rapid spread is unprecedented for monkeypox. Since the re-emergence of MPXV in Nigeria in 2017, there had only been two reported events of person-to-person transmission of MXPV outside of Africa: from a patient to a healthcare worker in the United Kingdom in 2018 (*4*) and a different patient to two family members in the United Kingdom in 2021 (*22*). Scientists have been scouring the limited publicly available sequence data to look for hints of viral evolution that could explain changes in viral proteins that would affect transmission. The MPXV genome from the USA_2021_MD case provides an opportunity to better focus genomic analysis, as this genome is the most similar to the predominant 2022 outbreak cluster out of all publicly available data. The 2021 Maryland case did not lead to any known secondary infections. There were only approximately 13 differences between the 2021 MD genome and the 2022 outbreak cluster, with seven causing changes in amino acid sequences. One of these was in the F13L (*Vaccinia virus* Copenhagen F13L homolog) coding sequence. F13L is the target of tecovirimat, the first antiviral agent specifically indicated for the treatment of smallpox. Functional studies are needed to determine if this mutation, shared by all sequences in the predominant 2022 outbreak cluster, affects the efficacy of tecovirimat against MPXV infection. Further examination of F13L and other mutations (Table 1, Table S1, Table S2) may be warranted to look for indicators of changes in viral behavior. However, it is currently unknown if the 2022 MPXV associated with most cases reported in Europe and the United States has evolved to be better at spreading between humans; other factors may also be at play, including behaviors involving close contact and failure to recognize or diagnose monkeypox to prevent spread.

Rapid diagnosis of monkeypox is critical for minimizing spread. Increased surveillance in the USA may have led to the identification of two monkeypox cases that are genetically unrelated to the predominant 2022 international outbreak variant. By careful comparison of Ct values from both the *Orthopoxvirus* OPX3 (*13*) and West African Monkeypox-specific (*14*) real-time PCR assays on validated platforms, laboratories can, in theory, differentiate between cases belonging to the 2022 outbreak lineage and cases from other lineages without sequencing. This observation of decreased sensitivity in the OPX3 assay is unrelated to the F510k cleared VAC1 assay (*23*) used for MPXV screening. Use of different commercial reagents, run parameters or PCR platform may result in different results; details for the OPX3 assay as performed in this study are provided in the methods.

Since 2017, West African MPXVs have all shared a striking number of G-to-A mutations that specifically occur in the context of the APOBEC3 motif 5’ GA-to-AA; others have observed this as well (*15*). This pattern was absent among older West African and Congo Basin MXPV lineages. APOBEC3 proteins are an important component of the vertebrate innate immune system that restrict the replication of exogenous viruses through cytosine-to-uracil deaminase activity (*16, 24*). APOBEC3 proteins act mainly on single stranded DNA and have been extensively studied in RNA viruses, including HIV (*17*) but can also act on DNA viruses (*25, 26*). *Vaccinia virus* (the prototypical orthopoxvirus) replication was not affected by APOBEC3 activity (*27*). Experiments of APOBEC3 activity in MPXV are needed to support and better understand the mutational patterns observed. The presence of APOBEC3 mutations in several branches of the West African MPXV lineage will cause errors in estimates of evolutionary rate and divergence time using many standard methods. Further studies are needed to understand APOBEC3 activity during MPXV infection and the host-viral interactions that led to the introduction of APOBEC3 changes in the West African MPXV lineage since 2017.

Indeed, mutations within the APOBEC3 motif were significantly enriched throughout the lineage that began with samples collected in 2017. No such pattern was observed among older West African or Congo Basin MPXV lineages, suggesting that something is different in the biology of the virus or the host specific interactions with the virus, facilitating this mutational pattern. Furthermore, each of 8 the mutations detected among current about outbreak samples studied here are 5’ GA-to-AA (Figure S6), indicating that this mutational bias is continuing. Poxviruses were not thought to be subject to APOBEC3 editing (*16, 27*). The fact that the enrichment can be observed at essentially all levels within the West African MPXV lineage since 2017 suggests it is a recurrent and dominant mutational effect in recent MPXV evolution.

## Methods: (Supplement)

### PCR testing

DNA was extracted from swabs collected from patient lesions using EZ-1 DNA tissue kit (Qiagen) followed by heat inactivation at 56°C for ≥ 1 hour. Monkeypox infection was confirmed by real-time PCR at CDC. West African-specific *Monkeypox virus* real-time PCR assay was performed as described in Li et al. (2010) (*14*). Orthopoxvirus OPX3 real-time PCR assay was performed as described (*13*), with the following changes: Each reaction (20 μL) contained 5 μL of template DNA, 0.5 μL of each primer (20 μM), 0.5 μL probe (10 μM) added to the 2X TaqMan Fast Advanced master mix (Applied Biosystems). Thermal cycling conditions for the ABI 7500 Fast Dx Real-Time PCR System (Applied Biosystems) were one cycle at 95°C for 20 seconds and 40 cycles at 95°C for 3 seconds and 60°C for 30 seconds.

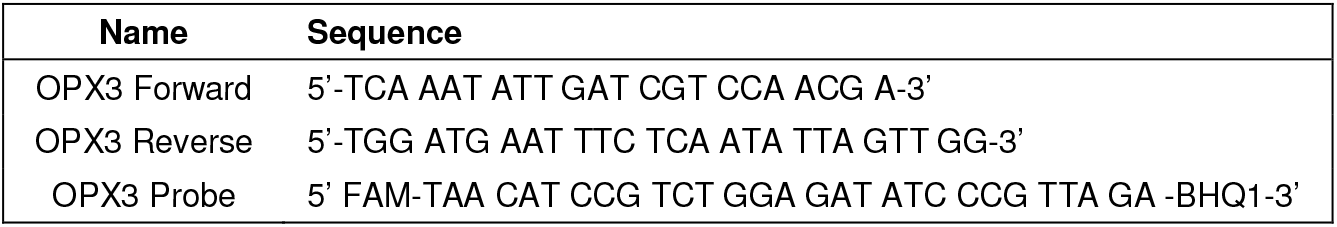

Table. Primer and probe sequences for the Orthopoxvirus OPX3 real-time PCR assay (called OPX in (*13*))

### Sequencing

ONT: Library preparation was performed on extracted DNA using Ligation Sequencing kit (Oxford Nanopore Technologies SQK-LSK-109) and sequenced on a MinION sequencer (MPXV_USA_2022_MA001, MPXV_USA_2021_MD, and MPXV_USA_2021_TX) or GridION sequencer (USA_2022_FL001, USA_2022_FL002, USA_2022_CA001, USA_2022_UT1) using a MIN109 R9.4.1 flow cell (Oxford Nanopore). For MPXV_USA_2022_MA001, data were combined from two sequencing runs from two independent swabs collected from two locations. For all other cases, data are from a single sample/swab. For the MinION run, basecalling was performed using guppy version with high accuracy and qscore filtering (for MinION runs) or was performed on GridION using high accuracy basecalling. Reads were mapped to MPXV West Africa Nigeria reference MT903343 (minimap2 2.16, https://github.com/lh3/minimap2) to remove human and non-MPXV reads. Consensus sequence was generating using samtools 1.7 and ivar (https://github.com/andersen-lab/ivar) with a 10x minimum read depth threshold and polished using medaka 1.0.1 (https://github.com/nanoporetech/medaka).

Illumina: Sequencing was conducted on the Illumina MiSeq platform using the Nextera XT sequencing kit according to the manufacturer’s instructions. De novo assembly was performed on reads that mapped to reference MT903343 (removing human and non-MPXV reads) using bwa mem using SPAdes. Consensus-based assembly was performed in CLC Genomics Workbench 22. Hybrid assembly with nanopore reads that mapped to MPXV MT903343 were generated using Unicylcer. Final genome assemblies were made by comparison of consensus-based and *de novo* assemblies. *De novo* nanopore assemblies and hybrid assembly were used to determine ITR and non-homopolymer repeat lengths. Illumina consensus sequences and contigs were used for error correction in homopolymer regions. Annotations were transferred from MPXV West Africa Nigeria reference MT903345, then locus_tags were re-named with the strain ID. Alignments used to make trees were performed using MAFFT v.7.450(*28*). Phylogenetic analysis was performed using BEAST v. 2.6.6. All sites containing gaps were removed prior to phylogenetic analysis, and all sites containing gaps or ambiguities were removed prior to haplotype network analysis. Due presence of ambiguities at the time of analysis, the following genomes were not included in analysis: Portugal PT0001, PT0002 (ON585030.1), PT0010 (ON585036.1) and 2022_Belgium. Still, many genomes contained Ns and may contain errors caused by low coverage or sequencing technology bias. Availability of complete, final genomes is expected to improve resolution of future analyses.

### APOBEC3 analysis

The sequence set we used started with foundational sequences from Mauldin et. al., a paper that described spread of MPXV beyond the African continent (*4*), adding additional sequences to create our baseline dataset of 84 MPXV-related sequences. Our initial alignment was generated using MAFFTv7 (*28*); ends were trimmed to exclude regions that were only sporadically sequenced, and the alignment further edited to resolve poorly aligned regions (generally these were in regions with differing numbers of direct repeats) and other inconsistencies. A maximum likelihood tree used for defining clades of interest and reconstructing ancestral states for subsequent APOBEC3 analysis in was generated using the HIV database web interface for IQ-tree (https://www.hiv.lanl.gov/content/sequence/IQTREE/iqtree.html) (*29*), with ModelFinder (*30*); the Bayesian information criterion (BIC) best fit model was K3Pu+F+I, and a mid-point root was used. All unique mutational events relative to the most common ancestor sequence within a given clade of interest were compressed onto a single sequence that represented unique mutations found within that clade, using the HIV database tool Highlighter (https://www.hiv.lanl.gov/content/sequence/HIGHLIGHT/highlighter_top.html). This compressed sequence was compared to the most recent common ancestor of the clade generated through IQ-tree (this comparison was used Figure 1B and Supp. Figures 4-6.

The statistical strategies for resolving if G-to-A mutations within an APOBEC3 motif context are enhanced are described in the HIV-database tool HYPERMUT (https://www.hiv.lanl.gov/content/sequence/HYPERMUT/hypermut.html) (*31*). The method makes pairwise comparisons between a reference sequence and each of a set of sequences aligned to that reference. It simply identifies and tallies all Gs that occur in the context of APOBEC3 motifs within a reference sequence and counts all G-to-A mutations that occur within the context of that motif in the sequence being compared; similarly, all Gs that are not in embedded in an APOBEC3 motif are tallied, and the number of G-to-A mutations that occur among those G’s are counted. A Fisher’s exact test is then used to determine if the G-to-A substitutions events are significantly enriched in the context of APOBEC3 motifs. Gaps and IUPAC ambiguity codes are excluded from the analysis. The original HYPERMUT strategy was developed for HIV-1 and so needed to be adapted for this application to enable simultaneously tracking G-to-A mutations in the forward and reverse complement strand. This was critical as what simply appears as C-to-T transition on the first strand can be embedded in the APOBEC3 motif in the reverse complement. HYPERMUT enables the inclusion of more complex APOBEC3 motifs, and we also explored the use of the 3-base motif pattern including a cytosine in the +2 positions (GAC-to-AAC, or GGC-to-AGC), as pattern this can partially inhibit APOBEC3 activity (*32*); as this strategy did not substantively change our conclusions, we present the simpler version here. Our method distinguishes between GA-to-AA and GG-to-AG motifs, and almost all of the substitutions we observed were in the GA to AA context, indicative of APOBEC3D and 3F activity, not APOBEC3G (*33*).

## Acknowledgements

We thank Brian Foley for providing a draft alignment of MPXV. BK was supported by the Laboratory Directed Research and Development program of Los Alamos National Laboratory under project number 20220660ER. We would like to thank Duncan MacCannell, Ketan Patel, Ying Tao, Sue Tong, Blake Cherney, and John Barnes from U.S. Centers for Disease Control and Prevention; Alexander Kim and Salimatu Lukula from the Maryland Department of Health Laboratory; Barbara Downes from Fairfax County, Virginia Department of Health, Catherine Brown, Larry Madoff, Mary DeMartino, Erika Buzby, Stephanie Ash, and Joshua Hall from the Massachusetts Department of Health; Juan Jaramillo from Dallas County LRN; Destiny Hairfield and Kimberly Stratton from Virginia Department of General Services - Division of Consolidated Laboratory Services.

---

Supplemental Table 1. List of mutations observed between USA_2022_MA001, USA_2021_MD and reference MT903344_2018_UK2.

Supplemental Table 2. List of mutations observed between USA_2022_FL001, USA_2022_VA001, USA_2021_TX and reference MT903344_2018_UK2.

Supplemental Table 3. List of average Ct values from *Orthopoxvirus* generic real-time PCR assay (OPX3) and West African-Specific Monkeypox virus real-time PCR assay. Averages are a result of three replicates.

Supplemental Table 4. List of sequences used in haplotype network analysis.

## Notes

### Competing Interest Statement

The authors have declared no competing interest.

