## Supplemental Figures 1 to 6 for "Multiple lineages of *Monkeypox virus* detected in the United States, 2021- 2022"

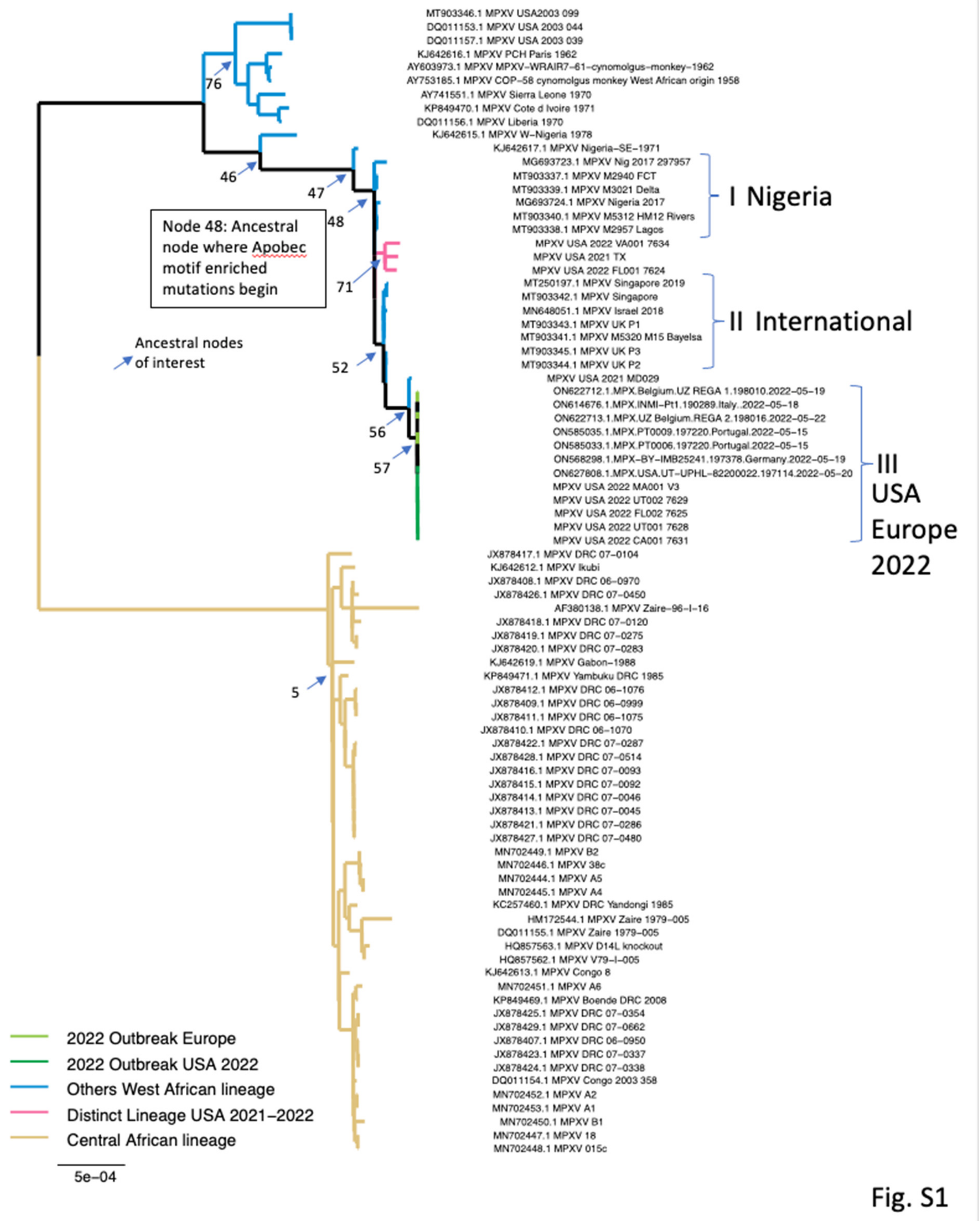


**Fig. S1. A detailed version of the ML tree shown in Fig. 1.** Ancestral nodes of interest are noted, and can be used to track the statistical exploration of APOBEC motif enrichment in Fig. S2. The sequence name of each of the taxa are shown. The full data for the Central African clade is provided.

**From Node 48:**

| | A B C D | E, F | |
|---|---|---|---|
| Node52 | [12 36570 1 30677] | p=0.004944, OR=10.07 | Branch A |
| Node71 | [14 36568 1 30677] | p=0.002755, OR=11.74 | Branch D |
| ON627808.1.MPX.USA.UT.197114.2022-05-20 | [58 36524 3 30675] | p=0.000000, OR=16.24 | |
| ON568298.1.MPX-.197378.Germany.2022-05-19 | [59 36523 3 30675] | p=0.000000, OR=16.52 | |
| ON585033.1.MPX.PT0006.197220.Portugal.2022-05-15 | [59 36523 3 30675] | p=0.000000, OR=16.52 | |
| ON585035.1.MPX.PT0009.197220.Portugal.2022-05-15 | [58 36524 3 30675] | p=0.000000, OR=16.24 | |
| MPXV_USA_2022_MA001_V3 | [58 36524 3 30675] | p=0.000000, OR=16.24 | Branch A+B+C |
| MPXV_USA_2022_CA001_7631 | [58 36524 3 30675] | p=0.000000, OR=16.24 | |
| MPXV_USA_2022_UT001_7628 | [59 36523 3 30675] | p=0.000000, OR=16.52 | |
| MPXV_USA_2022_FL002_7625 | [60 36522 3 30675] | p=0.000000, OR=16.8 | |
| MPXV_USA_2022_UT002_7629 | [58 36524 3 30675] | p=0.000000, OR=16.24 | |
| ON622712.1.MPX.Belgium.UZ_REGA_1.198010.2022-05-19 | [59 36523 3 30675] | p=0.000000, OR=16.52 | |
| ON622713.1.MPX.UZ_Belgium.REGA_2.198016.2022-05-22 | [60 36522 3 30675] | p=0.000000, OR=16.8 | |
| ON614676.1.MPX.INMI-Pt1.190289.Italy..2022-05-18 | [53 36529 3 30675] | p=0.000000, OR=14.84 | |
| MPXV_USA_2021_MD029 | [49 36533 4 30674] | p=0.000000, OR=10.29 | Sampled in 2021 |
| MT903338.1_MPXV_M2957_Lagos | [ 4 36578 0 30678] | p=0.130775, OR=inf | |
| MT903340.1_MPXV_M5312_HM12_Rivers | [ 5 36577 0 30678] | p=0.067325, OR=inf | |
| MG693724.1_MPXV_Nigeria_2017 | [ 7 36575 0 30678] | p=0.018181, OR=inf | Branch F |
| MT903339.1_MPXV_M3021_Delta | [ 4 36578 0 30678] | p=0.130775, OR=inf | |
| MT903337.1_MPXV_M2940_FCT | [ 3 36579 1 30677] | p=0.630746, OR=2.516 | |
| MG693723.1_MPXV_Nig_2017_297957 | [11 36571 0 30678] | p=0.001409, OR=inf | |
| MPXV_USA_2022_FL001_7624 | [28 36554 2 30676] | p=0.000008, OR=11.75 | |
| MPXV_USA_2021_TX | [32 36550 2 30676] | p=0.000001, OR=13.43 | |
| MPXV_USA_2022_VA001_7634 | [34 36548 2 30676] | p=0.000000, OR=14.27 | |

III. High similarity, all May 2022 MA001 is representative (ON627808.1 through ON614676.1)

I. Nigerian samples, only 2017 was significant, but across samples APOBEC motifs are enriched (MT903338.1 through MG693723.1)

Second US introduction (MPXV_USA_2022_FL001_7624, MPXV_USA_2021_TX, MPXV_USA_2022_VA001_7634)

**From Node 71: Second US clade**

| | A B C D | E, F | |
|---|---|---|---|
| MPXV_USA_2022_FL001_7624 | [14 36554 1 30676] | p=0.002755, OR=11.75 | |
| MPXV_USA_2021_TX | [18 36550 1 30676] | p=0.000248, OR=15.11 | Branch E |
| MPXV_USA_2022_VA001_7634 | [20 36548 1 30676] | p=0.000074, OR=16.79 | |

**From Node 46:**

| | A B C D | E, F |
|---|---|---|
| KJ642615.1_MPXV_W-Nigeria_1978 | [12 36596 20 30692] | p=0.074390, OR=0.5032 |

**From Node 47:**

| | A B C D | E, F |
|---|---|---|
| Node48 | [16 36579 8 30675] | p=0.305706, OR=1.677 |

**From Node 52:**

| | A B C D | E, F | |
|---|---|---|---|
| Node56 | [35 36535 2 30675] | p=0.000000, OR=14.69 | Branch B |
| MPXV_USA_2022_MA001_V3 | [46 36524 2 30675] | p=0.000000, OR=19.32 | |
| MPXV_USA_2021_MD029 | [37 36533 3 30674] | p=0.000000, OR=10.36 | |
| MT903344.1_MPXV_UK_P2 | [ 3 36567 0 30677] | p=0.255748, OR=inf | |
| MT903345.1_MPXV_UK_P3 | [ 3 36567 0 30677] | p=0.255748, OR=inf | |
| MT903341.1_MPXV_M5320_M15_Bayelsa | [ 2 36568 0 30677] | p=0.503832, OR=inf | |
| MT903343.1_MPXV_UK_P1 | [ 2 36568 1 30676] | p=1.000000, OR=1.678 | |
| MN648051.1_MPXV_Israel_2018 | [ 4 36566 0 30677] | p=0.130756, OR=inf | |
| MT903342.1_MPXV_Singapore | [ 4 36566 0 30677] | p=0.130756, OR=inf | |
| MT250197.1_MPXV_Singapore_2019 | [ 4 36566 0 30677] | p=0.130756, OR=inf | |

!!. Recent global samples, few mutations in any one but across samples APOBEC motifs are enriched (MT903344.1 through MT250197.1)

**From Node 56:**

| | A B C D | E, F | |
|---|---|---|---|
| Node57 | [11 36524 0 30675] | p=0.001403, OR=inf | Branch C |
| MPXV_USA_2022_MA001_V3 | [11 36524 0 30675] | p=0.001403, OR=inf | |

Fig. S2

**Figure S2. Statistical analysis of the enrichment of APOBEC-motif substitution in the recent West African lineage data.** The ML estimated ancestral sequence for different nodes in the tree are compared taxa sequences at leaves or to relevant intermediated ancestral nodes in each group. Columns: (A) The number of G-> A changes in an APOBEC context, GR -> AR. (B) The number of GR motifs in the reference strain where the initial G remains unchanged. (C) The number of G-> A changes not in an APOBEC context, GY -> AY. (D) The total number of G's in the reference strain in a GY context where the G remained unchanged. (E) A two-sided Fisher's exact p-value of the contingency table based on columns A-D. (F) The odds ratio (OR) of (A/(A+B))/(C/C+D), numbers greater than 1 indicating the level of relative enrichment of G-> A changes in an APOBEC context. Notes on the right indicate relevant parts of the tree show in Slide 2 and 3. Blue highlights indicate branches in the tree where G-to-A changes in APOBEC motifs are statistically enriched. There was no evidence of G->A mutations embedded in APOBEC3 motifs within the older West African or the Central African lineages (details of this analysis are not shown).

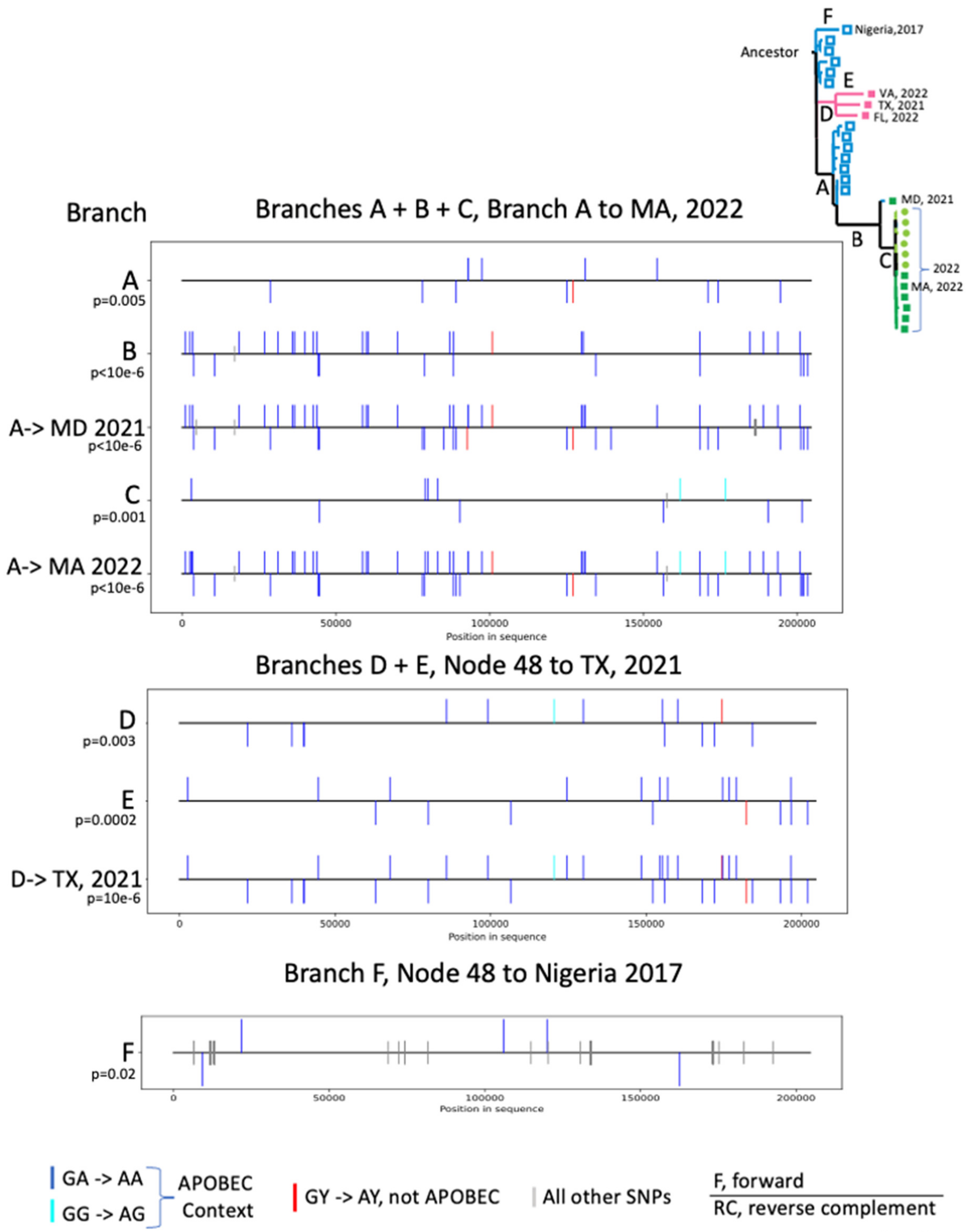


**Fig. S3. All SNP mutations on branches with significant APOBEC motif enrichment.** This illustration indicates where all SNPs are located in the genome that arose in different branches in the phylogeny (labeled in Fig. 1 and Fig. S2), featuring those that were significantly enriched for APOBEC-context G-to-A substitutions. GA-to-AA patterns are indicated in blue; they are overwhelmingly the most commonly found substitutions on both the forward and reverse strand. The p-values supporting the APOBEC motif enrichment are shown on the left, and details regarding how these statistics were calculated are provided in Fig. S2. The branches A+B+C leading to the May 2022 outbreak (represented by MA, 2022), and the branches D+E leading to the second introduction variant TX, 2021, are each significantly enriched for G-to-A in an APOBEC motif. Branch F, leading to the Nigerian 2017 sequence was also enriched for GA-to-AA substitutions relative to G-to-A outside of an APOBEC motif, but more subtly, as other kinds of SNPs are more common.

### Cluster I: comparison to most recent common ancestor, node 52

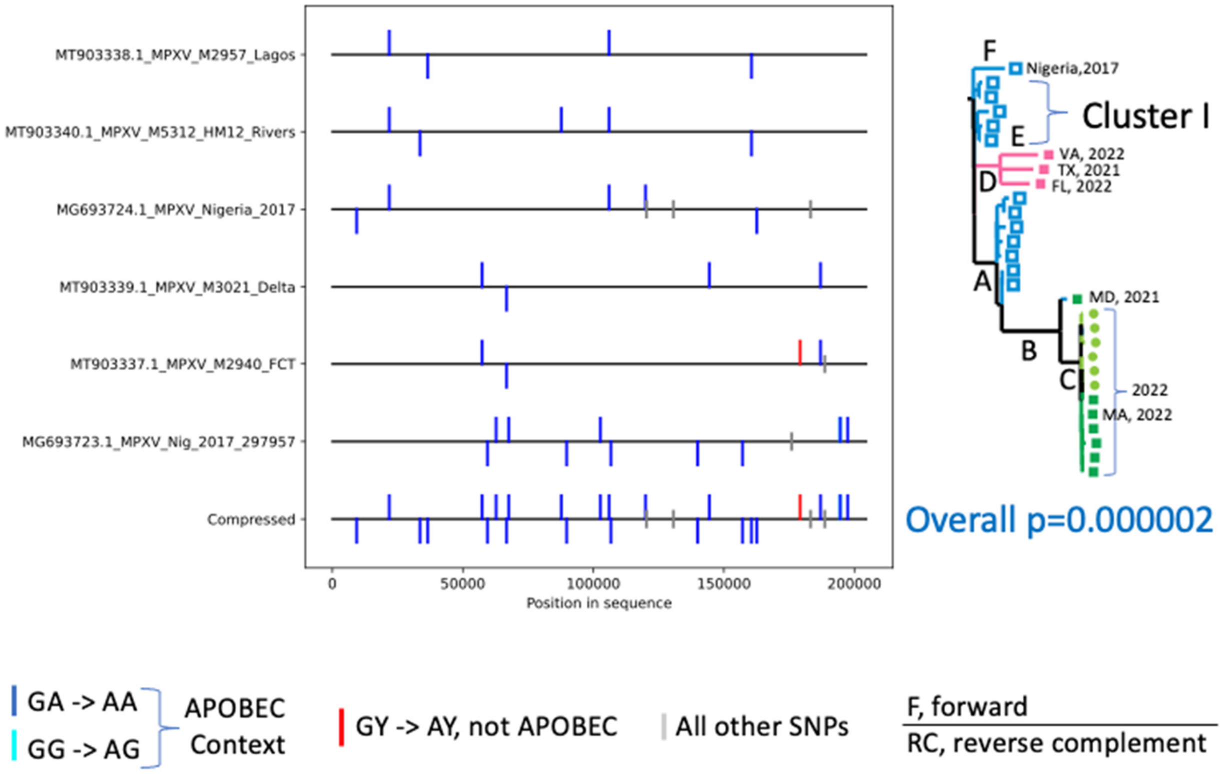


#### Counts of mutations and statistics

| sequence-name | other-mutations | [contingency-table] | p-value | odds-ratio |
|---|---|---|---|---|
| MT903338.1_MPXV_M2957_Lagos | 0 | [ 4 36578 0 30678] | p=0.130775, | OR=inf |
| MT903340.1_MPXV_M5312_HM12_Rivers | 0 | [ 5 36577 0 30678] | p=0.067325, | OR=inf |
| MG693724.1_MPXV_Nigeria_2017 | 3 | [ 7 36575 0 30678] | p=0.018181, | OR=inf |
| MT903339.1_MPXV_M3021_Delta | 0 | [ 4 36578 0 30678] | p=0.130775, | OR=inf |
| MT903337.1_MPXV_M2940_FCT | 1 | [ 3 36579 1 30677] | p=0.630746, | OR=2.516 |
| MG693723.1_MPXV_Nig_2017_297957 | 1 | [11 36571 0 30678] | p=0.001409, | OR=inf |
| Compressed | 4 | [26 36556 1 30677] | p=0.000002, | OR=21.82 |

Fig. S4

**Fig. S4. SNP mutations that arose within the highly related set of sequences from Nigeria found in cluster I were almost all G-to-A substitutions embedded in a GA-to-AA APOBEC motif.** There were 26 G-to-A mutations within an APOBEC motif, only 2 G-to-As that were not in an APOBEC motif, and only 4 mutations that were not to G-A on either the forward or reverse complement strand. Though only the Nigeria 2017 sequence showed modest enrichment for APOBEC motif mutations when considered in isolation, the across the full cluster such mutations were highly significantly enriched.

### Cluster II: comparison to most recent common ancestor, node 52

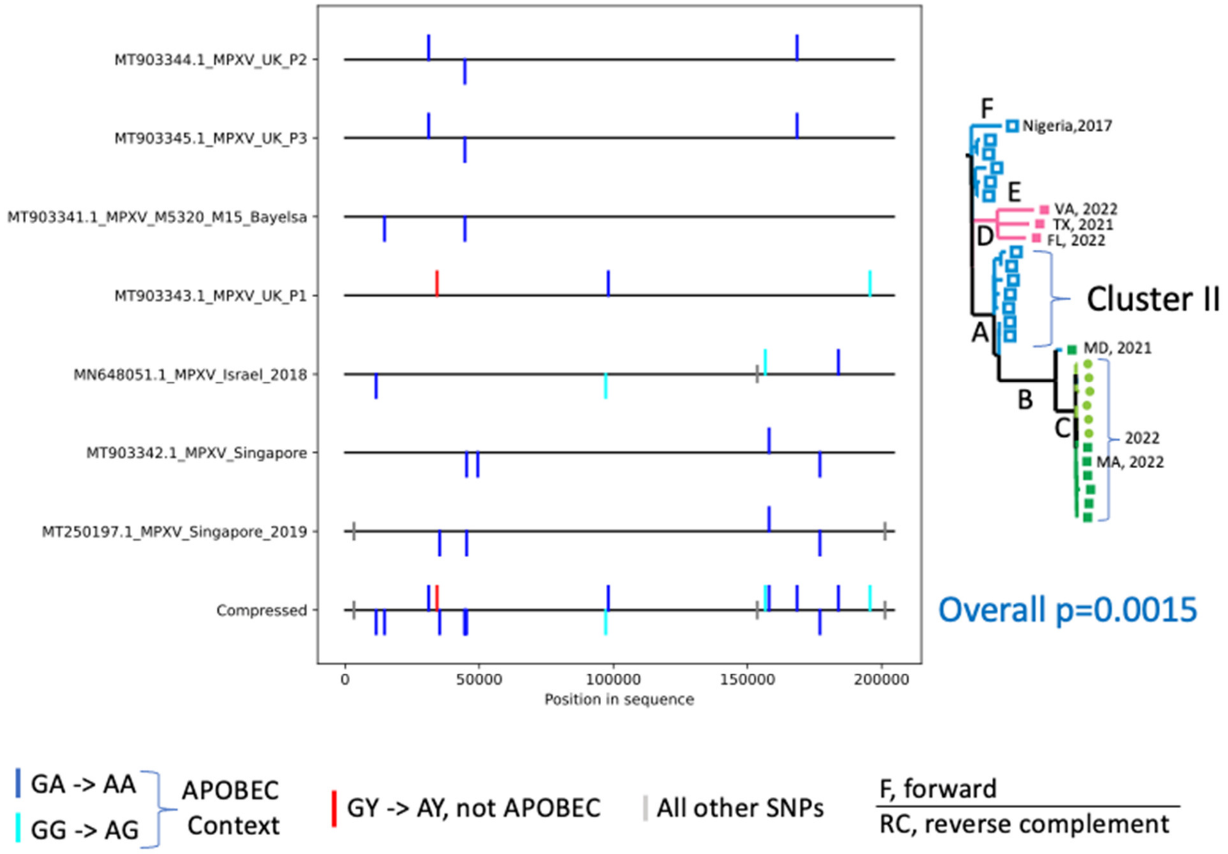


#### Counts of mutations and statistics

| sequence-name | other-mutations | [contingency-table] | p-value | odds-ratio |
|---|---|---|---|---|
| MT903344.1_MPXV_UK_P2 | 0 | [ 3 36567 0 30677] | p=0.255748, | OR=inf |
| MT903345.1_MPXV_UK_P3 | 0 | [ 3 36567 0 30677] | p=0.255748, | OR=inf |
| MT903341.1_MPXV_M5320_M15_Bayelsa | 0 | [ 2 36568 0 30677] | p=0.503832, | OR=inf |
| MT903343.1_MPXV_UK_P1 | 0 | [ 2 36568 1 30676] | p=1.000000, | OR=1.678 |
| MN648051.1_MPXV_Israel_2018 | 1 | [ 4 36566 0 30677] | p=0.130756, | OR=inf |
| MT903342.1_MPXV_Singapore | 0 | [ 4 36566 0 30677] | p=0.130756, | OR=inf |
| MT250197.1_MPXV_Singapore_2019 | 2 | [ 4 36566 0 30677] | p=0.130756, | OR=inf |
| Compressed | 3 | [ 15 36555 1 30676] | p=0.001512, | OR=12.59 |

Fig. S5

**Fig. S5. SNP mutations that arose within the highly related set of sequences in cluster II isolated when MPXV began started to be isolated outside of Africa in 2017.** Mutations within cluster II were almost all G-to-A substitutions embedded in a GA-to-AA APOBEC motif. There were 15 G-to-A mutations within an APOBEC motif, only 1 G-to-As that was not in an APOBEC motif, and only 3 mutations that were not to G-A on either the forward or reverse complement strand. No single sequence showed enrichment for APOBEC motif mutations when considered in isolation, but across the full cluster of such mutations were significantly enriched.

### Cluster III: comparison to most recent common ancestor Node 57

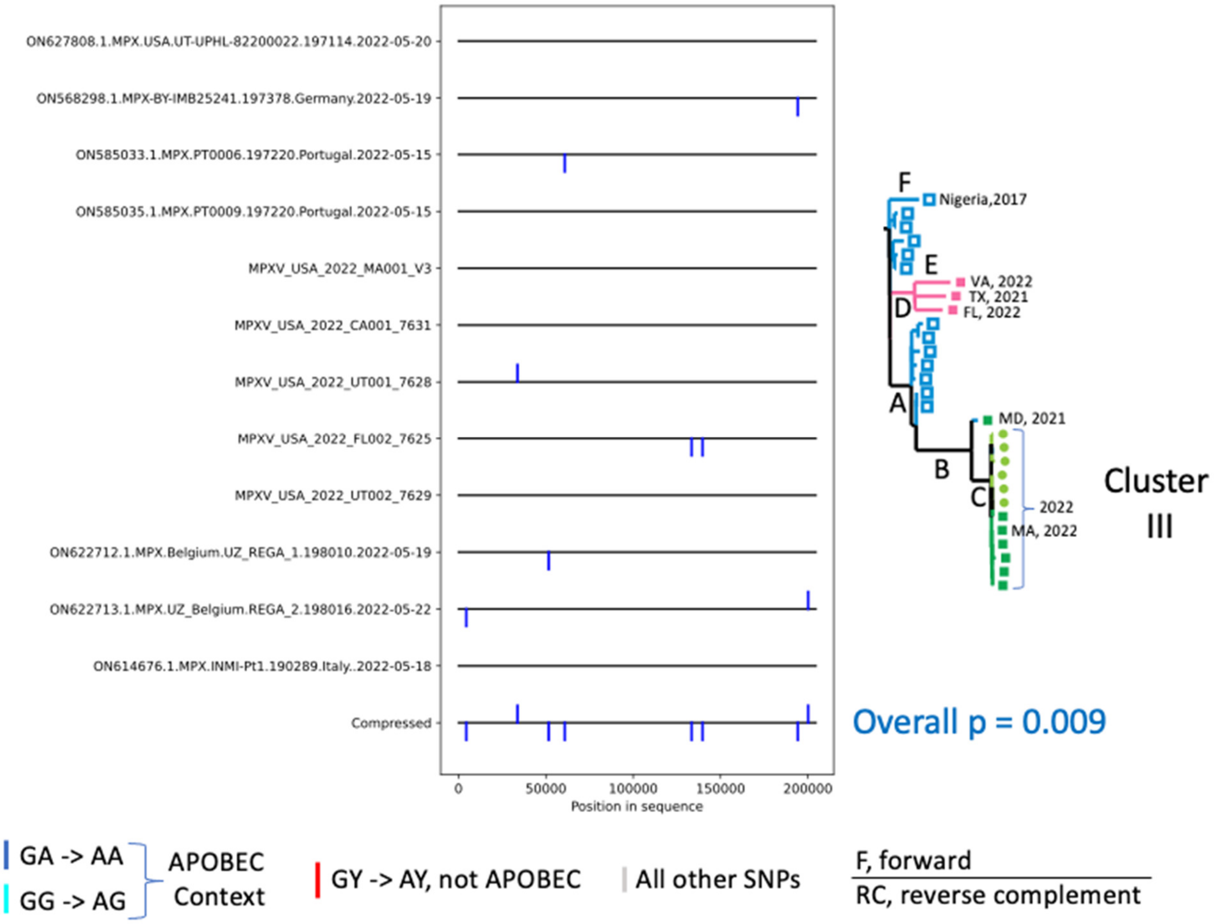


#### Counts of mutations and statistics

| sequence-name | other-mutations | [contingency-table] | p-value | odds-ratio |
|---|---|---|---|---|
| ON627808.1.MPX.USA.UT-UPHL-82200022.197114.2022-05-20 | 0 | [ 0 36525 0 30675] | p=1.000000, | OR=nan |
| ON568298.1.MPX-BY-IMB25241.197378.Germany.2022-05-19 | 0 | [ 1 36524 0 30675] | p=1.000000, | OR=inf |
| ON585033.1.MPX.PT0006.197220.Portugal.2022-05-15 | 0 | [ 1 36524 0 30675] | p=1.000000, | OR=inf |
| ON585035.1.MPX.PT0009.197220.Portugal.2022-05-15 | 0 | [ 0 36525 0 30675] | p=1.000000, | OR=nan |
| MPXV_USA_2022_MA001_V3 | 0 | [ 0 36525 0 30675] | p=1.000000, | OR=nan |
| MPXV_USA_2022_CA001_7631 | 0 | [ 0 36525 0 30675] | p=1.000000, | OR=nan |
| MPXV_USA_2022_UT001_7628 | 0 | [ 1 36524 0 30675] | p=1.000000, | OR=inf |
| MPXV_USA_2022_FL002_7625 | 0 | [ 2 36523 0 30675] | p=0.503782, | OR=inf |
| MPXV_USA_2022_UT002_7629 | 0 | [ 0 36525 0 30675] | p=1.000000, | OR=nan |
| ON622712.1.MPX.Belgium.UZ_REGA_1.198010.2022-05-19 | 0 | [ 1 36524 0 30675] | p=1.000000, | OR=inf |
| ON622713.1.MPX.UZ_Belgium.REGA_2.198016.2022-05-22 | 0 | [ 2 36523 0 30675] | p=0.503782, | OR=inf |
| ON614676.1.MPX.INMI-Pt1.190289.Italy..2022-05-18 | 0 | [ 0 36525 0 30675] | p=1.000000, | OR=nan |
| Compressed | 0 | [ 8 36517 0 30675] | p=0.009498, | OR=inf |

**Fig. S6. SNP mutations that arose within the highly related set of sequences in cluster III isolated from the May 2022 outbreak, sampled in Europe and the USA.** Mutations within cluster III were most often G-to-A substitutions embedded in a GA-to-AA APOBEC motif. Among the recent outbreak sequences, there are 8 GA-to-AA mutations within an APOBEC motif, no G-to-As not in an APOBEC motif, and no mutations that were not to G-A on either the forward or reverse complement strand. No single sequence showed enrichment for APOBEC motif mutations when considered in isolation, but across the full cluster such mutations were highly significantly enriched.
